# Algorithmic complexity of EEG for prognosis of neurodegeneration in idiopathic rapid eye movement behavior disorder (RBD)

**DOI:** 10.1101/200543

**Authors:** Giulio Ruffini, David Ibañez, Eleni Kroupi, Jean-François Gagnon, Jacques Montplaisir, Ronald B. Postuma, Marta Castellano, Aureli Soria-Frisch

**Affiliations:** Starlab Barcelona, Barcelona; Centre for Advanced Research in Sleep Medicine, Hôpital du Sacré- Coeur de Montréal, Montreal, Canada

## Abstract

**Objective:** Idiopathic REM sleep behavior disorder (RBD) is a serious risk factor for neurodegenerative processes such as Parkinson’s disease (PD). We investigate the use of EEG algorithmic complexity derived metrics for its prognosis.

**Methods:** We analyzed resting state EEG data collected from 114 idiopathic RBD patients and 83 healthy controls in a longitudinal study forming a cohort in which several RBD patients developed PD or dementia with Lewy bodies. Multichannel data from *∼*5 minute recordings was converted to spectrograms and their algorithmic complexity estimated using Lempel-Ziv-Welch compression (LZW).

**Results:** Complexity measures and entropy rate displayed statistically significant differences between groups. Results are compared to those using the ratio of slow to fast frequency power, which they are seen to complement by displaying increased sensitivity even when using a few EEG channels.

**Conclusions:** Poor prognosis in RBD appears to be associated with decreased complexity of EEG spectrograms stemming in part from frequency power imbalances and cross-frequency amplitude coupling.

**Significance:** Algorithmic complexity metrics provide a robust, powerful and complementary way to quantify the dynamics of EEG signals in RBD with links to emerging theories of brain function stemming from algorithmic information theory.

**Index Terms:** Biomarkers, EEG, LZW, PD, LBD

## I. INTRODUCTION

REM Behavior Disorder (RBD) is a serious risk factor for neurodegenerative diseases such as Parkinson’s disease (PD). RBD is a parasomnia characterized by vivid dreaming and dream-enacting behaviors associated with REM sleep without muscle atonia [1]. Idiopathic RBD occurs in the absence of any neurological disease or other identified cause, is male-predominant and its clinical course is generally chronic progressive [2]. Several longitudinal studies conducted in sleep centers have shown that most patients diagnosed with the idiopathic form of RBD will eventually be diagnosed with a neurological disorder such as Parkinson disease (PD) and dementia with Lewy bodies (DLB) [1]–[3]. In essence, idiopathic RBD has been suggested as a prodromal factor of the synucleinopathies PD, DLB and less frequently multiple system atrophy (MSA) [1].

RBD has an estimated prevalence of 15-60% in PD and has been proposed to define a subtype of PD with relatively poor prognosis, reflecting a brainstem-dominant route of pathology progression (v. [4] and references therein) with a higher risk for dementia or hallucinations. PD with RBD is characterized by more profound and extensive pathology—not limited to the brainstem—,with higher synuclein deposition in both cortical and sub-cortical regions.

The human brain can be modeled as a highly dimensional complex dynamical system which instantiates electrochemical communication and computation. Electroencephalographic (EEG) and magnetoencephalographic (MEG) signals contain rich information associated with these processes, and are accessible non-invasively. To a large extent, progress in the analysis of such signals has been driven by the study of classical temporal and spectral features in electrode space, and applied to the study the human brain in both health and disease. For example, the “slowing down” of EEG is known to characterize neurodegenerative diseases [5], [6], and the slow to fast ratio (the ratio of power in delta and theta bands to alpha and beta) has shown good discriminatory sensitivity [6], [7]. However, brain activity measurements exhibit nonlinear dynamics and non-stationarity, limiting the usefulness of classical, linear approaches and calling for the use of novel methods capable of exploiting underlying spatiotemporal hierarchical structures. Deep learning techniques in particular and neural networks in general are bio-inspired by neural structure and function—the same biological systems generating the electric signals we aim to decode—and should be well suited for the task. In past work, for example, we studied a particular class of recurrent neural networks called Echo State Networks (ESNs) combining the power of networks for classification of temporal patterns and ease of training, reasoning that as they implement non-linear dynamics with memory they may be ideally poised for the classification of complex EEG time series data. Starting from a dataset of recordings of resting state EEG from idiopathic RBD patients who later converted to PD and from healthy controls (HC), we showed that using such recurrent networks using temporal series of EEG power led to a prognosis classification accuracy of 85% in the binary, balanced classification problem [8].

With the goal of developing metrics that capture non-linear dynamics from EEG signals, here we explore a different feature extracted from EEG data—upper bounds on algorithmic complexity. Algorithmic information is formalized by the mathematical concept of algorithmic complexity or Kolmogorov complexity (*𝒦*), co-discovered during the second half of the 20th century by Solomonoff, Kolmogorov and Chaitin. We recall its definition: *the Kolmogorov complexity of a string is the length of the shortest program capable of generating it.* More precisely, let 𝒰 be a universal computer (a Turing Machine), and let *p* be a program. Then the Kolmogorov or algorithmic complexity of a string *x* with respect to 𝒰 is given by *𝒦*_𝒰(*x*)_ = min_*p*: 𝒰(*p*)=*x*_ *l*(*p*), i.e., the length *l*(*p*) of the shortest program that prints the string *x* and then halts (see e.g., [9], [10]). Crucially, although the precise length of this program depends on the programming language used, it does so only up to a string-independent constant. Algorithmic information is notoriously difficult to estimate. Here we rely on the notion of complexity as measured from Lempel-Ziv-Welch compression (LZW) or entropy rate, two closely related metrics that provide an upper bound to algorithmic complexity.

As discussed in [11] and references therein, the healthy brain generates apparently complex (entropic) data. Complexity should be associated to cognitive health and conscious state, and complexity appears to decrease with age (see, e.g., [12] and references therein). While a brain capable of universal computation may produce many different types of patterns— from simple to highly entropic—a healthy brain engaging in modeling, prediction and interaction with the world will produce complex-looking, highly entropic data which can presumably be measured from behavior or directly in brain activity. Entropy and LZW represent direct measures of apparent complexity and can be applied to, e.g., electrophysiological or metabolic brain data [13]–[17]. We hypothesize here that global apparent EEG algorithmic complexity or entropy rate in our dataset will decrease with worsening neurodegenerative disease prognosis or progression, in a similar manner to that already reported in the PD literature [18] and in the case of Alzheimer’s disease [19]–[21]. As a starting point, we further assume that algorithmically relevant aspects in EEG data are present in compositional features in the time-frequency spectral amplitude representation.

## II. METHODS

### A. Participants

Idiopathic RBD patients (called henceforth RBD for data analysis labeling) and healthy controls were recruited at the Center for Advanced Research in Sleep Medicine of the Hôpital du Sacrè-Cœur de Montral. All patients with full EEG montage for resting-state EEG recording at baseline and with at least one follow-up examination after the baseline visit were included in the study. The first valid EEG for each patient enrolled in the study was considered baseline. Participants also underwent a complete neurological examination by a neurologist specialized in movement disorders and a cognitive assessment by a neuropsychologist. No controls reported abnormal motor activity during sleep or showed cognitive impairment on neuropsychological testing. The protocol was approved by the hospital’s ethics committee, and all participants gave their written informed consent to participate. For mode details, the reader is directed to [6].

### B. EEG dataset

As with previous work [22], the raw data in this study consisted of resting-state EEG collected from awake patients and healthy controls using a subset of 14 scalp electrodes (C3, C4, F3, F4, F7, F8, O1, O2, P3, P4, P7, P8, T7, T8). The recording protocol consisted of conditions with periods of “eyes open” of variable duration (approximately two minutes) followed by periods of “eyes closed” in which patients were not given any particular task. Resting EEG signals were digitized with 16-bit resolution at a sampling rate of 256 S/s. The amplification device used a hardware band pass filter between 0.3 and 100 Hz and a line-noise notch filter at 60 Hz. All recordings were referenced to linked ears.

After quality check with rejection of subjects with insufficient data or subjects with abnormal cognition at the time of acquisition, the dataset (of initially 213 subjects) consisted of eyes-closed resting EEG data from a total of 114 patients who were diagnosed with idiopathic RBD at the time of acquisition and 83 healthy controls without sleep complaints in which RBD was excluded. EEG data was collected in every patient at baseline, i.e., when they were still diagnosed with idiopathic RBD. After 1–10 years of clinical followup, 19 patients developed Parkinson disease (PD), 12 Lewy body dementia (DLB) and the remaining 83 ones remained idiopathic RBD. Summary demographic data are provided in Table I. Group averaged power spectral density plots for each group are provided in Figure 1.

**TABLE I.** Baseline sociodemographic and clinical data for the four groups.

| Group / data | HC | RBD | PD | DLB |
| --- | --- | --- | --- | --- |
| N (F/M) | 83 (30/53) | 83 (18/65) | 19 (5/14) | 12 (2/10) |
| <age> | 66 ± 9 | 67 ± 7 | 67 ± 7 | 71 ± 6 |
| <follow-up> | 1 ± 1 | 4 ± 3 | 4 ± 2 | 4 ± 2 |

Number of subjects per group with gender breakdown (female/male), mean and standard deviation for age and follow-up time (in years). Acronyms: HC: healthy controls; RBD: idiopathic RBDs who remained idiopathic RBD on follow-up; PD: idiopathic RBD who converted to PD on follow-up; DLB: idiopathic RBD who converted to LBD on follow-up.

**Fig. 1.**
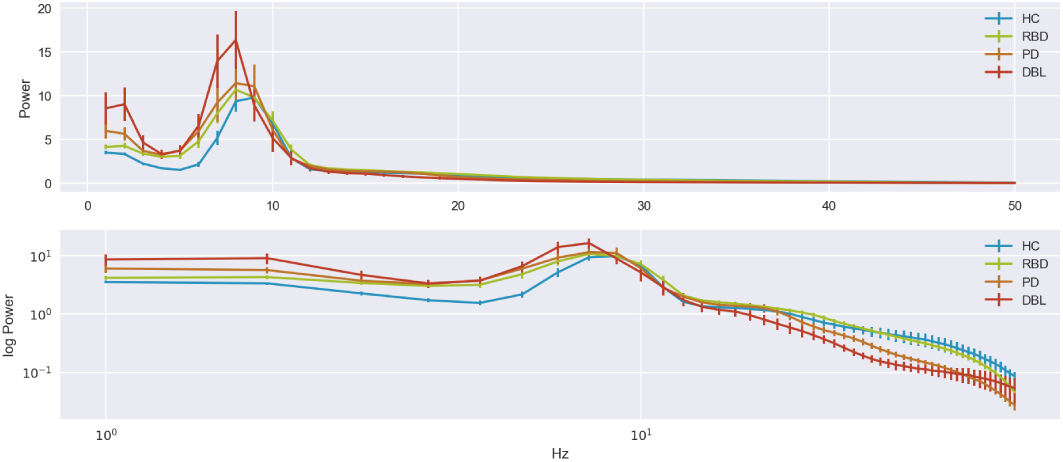
Group PSDs. Average power Spectral Density (all channels) for each group with standard error of the mean (SEM), displaying the characteristic “slowing” of EEG with more power at low frequencies.

### C. Data preparation

First, the EEG dataset from each subject was converted into a spectrogram stack of “frames” (time frequency bidimensional arrays) (see Figure 2), with each stack representing about 3 minutes of eyes-closed resting EEG. To generate it, EEG data for each channel was processed using Fourier analysis (FFT) after detrending one-second data blocks with a Hann window (50% overlap FFT with resolution is 1 Hz in the range 1–50Hz). Each spectrogram frame was generated from 20 second long artifact free sequences for each of the 14 EEG channels using a sliding window of one second between frames. The resulting data frames are thus “tensors” of the form [channels (14)] x [FFT bins (50)] x [Time bins (39)], and the full stack is four dimensional, with the fourth dimension indexing frame epoch.

**Fig. 2.**
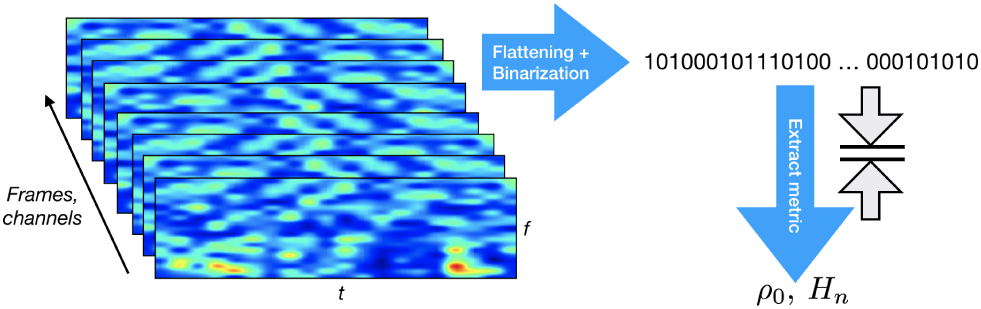
Computation of description length and entropy metrics from subject spectrogram frame stack. To compute a global measure of complexity for each subject, the frame stack *S*(*t, f, ch, F*) is flattened and binarized prior compression or entropy computation, where *t* denotes time in each spectrogram, *f* frequency, *ch* channel, and *F* frame number.

Using the spectrogram frame stack dataset we carried out an analysis of the complexity for each subject, as we discuss next.

### D. Lempel-Ziv-Welch complexity and entropy rate

As discussed above, in order to provide an upper bound to description length or algorithmic complexity we use LZW compression. The LZW algorithm seeks increasingly long reappearing patterns in the data, and compresses information by reusing them: instead of rewriting a previously seen sequence, it will refer to the identifier of the one seen last [23]. After applying LZW to a string, we are provided with a set of words (or phrases, as they are sometimes called) *c*(*n*) that form a dictionary. The length of the compressed string will, in general, be *l*_*LZW*_ *≲ n*. We note that LZW can be used to obtain an upper bound on algorithmic complexity, but, given its reduced programming repertoire (LZW is the Kolmogorov complexity computed with a limited set of programs that only allow copy and insertion in strings [24]), it will fail to compress random-looking data generated by simple, but highly recursive programs, e.g., the sequence of binary digits of *π*, with more sophisticated regularities than sequence repetition (deep programs [8]). As an example of how LZW or entropy rate are limited tools for compression, and therefore coarse approximations to algorithmic complexity, consider a string and its bit-flipped version, or a “time-reversed” version in a file with the ordering of symbols temporally inverted, or a string and “time-dilated” string where each symbol is repeated, say twice. Such simple algorithmic manipulations will not be detected and exploited by LZW. Despite these limitations, such compression algorithms can be useful, as we shall see below. Let *H*_0_(*p*) = −*p* log *p* − (1 − *p*) log(1 − *p*) denote the univariate or zero order entropy, with *p* the probability of a Bernoulli (binary) process (Markov chain of order zero). By the entropy rate of the stochastic process {*X*_*i*_}, we will mean

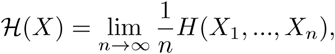

when this limit exists, with *H* denoting the usual multivariate entropy of *X, H*(*X*) = − *E*_*X*_ [log(*P* (*X*)]. We note that entropy rate of a stochastic processes is non-increasing as a function of order, that is, 0*≤ ℋ ≤. ≤H*_*q*_*≤ … ≤H*_0_*≤* 1. A fundamental relation is that description length computed from LZW is closely related to entropy rate, that is,

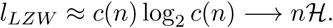

(see [9], [25]). Thus, the description length of the sequence encoded by LZW can be approximated by the number of phrases times the number of bits needed to identify a seen phrase. In order to apply LZW we need first to digitize the data. Here we binarize the data to reduce the length of strings needed for LZW to stabilize. A reasonable strategy in analysis of algorithmic information of EEG data is to preserve as much information as possible in the resulting transformed string. In this sense, using methods that maximize the entropy of the resulting series are recommended, such as using the median for thresholding (this is guaranteed to result in *H*_0_ = 1 and makes the result independent of overall scale).

Two associated metrics are commonly used in the field: *c*(*n*) and *l*_*LZW*_. Of the two, the latter is more closely related to Kolmogorov complexity or description length. Both contain similar information (in fact one is a monotonic function of the other). A natural manner to normalize this metric is to divide description length by the original string length *n, ρ*_0_ = *l*_*LZW*_ */n→ ℋ*, with units of bits per character. This is the LZW compression ratio metric we will use here. For details and code used to compute LZW complexity see [25].

### E. Data flattening and binarization

LZW and entropy rate can be sensitive to inherent choices in data flattening (that is, how the multidimensional 4D arrays are flattened into a one dimensional string). Here we have focused on the search for temporal patterns. For this purpose, the stack data tensor for each subject was converted to a one-dimensional array maintaining temporal adjacency in each frame (with flattening in this order: time in each frame, epoch, channel, frequency). The spectrogram hypercube flattened arrays for each subject were then binarized using the median as a threshold. We note here that binarization will produce sequences with information related to burst or “bump” events in each band. Moreover, these sequences are independent of overall scale of the data, since we used the median for binarization. We computed *ρ*_0_ and the entropy rate for orders 0 (*H*_0_) to 5 (*H*_5_, there are 2^6^ =64 cases or bins in the associated histogram, which is a reasonable number given the length of the strings analyzed), and computed the log-log line fit slope and clustering coefficient of MAI derived graphs using *μ*_0_ as an additional metric. Since these metrics can be sensitive to data string length [24], for the computational of global complexity we chose a minimal string length and fixed it for all subjects data prior compression (using the first N bits of the flattened spectrogram frame stacks, with *N* = 4.6 × 10^6^, or about 169 twenty-second epochs, as a reasonable tradeoff between number of subjects and data quantity per subject). In an analysis variant we carried out the complexity analysis frame by frame, producing an estimate for each frame and then an average of frame complexity across epochs per subject. In this case, we could use all the available data per subject, since string length equality in LZW was then guaranteed regardless of number of epochs used for averaging.

### F. Mutual information

A measure such as *ρ*_0_ applied to multi-channel, multi-epoch spectrograms reflects any detectable regularities across time, channels and frequencies in the frame stacks, and exploit, in that sense, the integration of multichannel, multifrequency EEG data. We can apply it in a global sense by compressing the entire dataset, that is, all channels and frequencies and epochs forming a single input data string. Or, we can study different data subsets to further dissect the driving elements of integration (compression) of the global dataset. We can ask, for example, how much is to be gained in the compression of a spectrogram stack by feeding all the stack (channels and frequencies) into the compression engine, compared to what can be achieved compressing, for example, each frequency independently and then adding the results. The more “integrated” the data is, the more is to be gained by global compression. If there is mutual algorithmic information across frequency bands, for, example, LZW may be able to use it, making a better job than compressing one channel at the time. In this vein, we can define several compression “integration” measures, depending along what axis we want to segment spectrogram stack data, for example channel or frequency.

To do this we start from the *mutual algorithmic information (MAI)* between two strings [26], the algorithmic analog of Shannon mutual information. Let 𝒦 (*x*) denote the description length of a string *x*, which we approximate by the LZW description length. The MAI between two strings is then

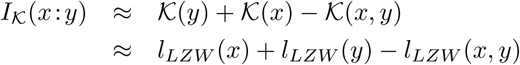

where *K*(*x, y*) denotes the complexity of the concatenated strings, and where we again approximate Kolmogorov complexity by LZW description length (see [11], annotated version). We now define the mutual algorithmic information coefficient between to strings to be

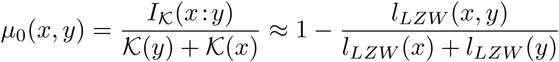

This metric is closely related to the Normalized Compression Distance [27], [28]. Using it we can assess the MAI between frequency bands, for example, and build an adjacency matrix or graph defined by a set of nodes (e.g., frequencies) and edges or links (MAI between nodes using a threshold). In this way, we associate a mutual information graph with each subject frame stack. We can then analyze these graphs using standard metrics such as average degree or clustering index *C* [29], [30]. Such analysis can also help analyze how compression of the data is actually taking place, as we discuss below.

## III. RESULTS

Entropy rate estimates as a function of order for each group are shown in Figure 3, displaying clear differences across two super-groups (HC+RBD vs PD+DLB) and also a flattening out with increasing order. LZW complexity metrics per subject are shown, sorted, in Figure 4 together with the slow-to-fast ratio (further discussed below), and summarized in Figure 5. In terms of statistical performance, the related complexity related metrics we tested (*ρ*_0_, entropy rate and log-log entropy rate slope) produced similar results. Statistical results for the main metrics are summarized in Table II (mean with standard error of the mean and four-way Kruskal-Wallis test). Since *H*_5_ entropy rate or power law slope led to very similar results to those using *ρ*_0_ (which is much faster to compute), we will drop them in what follows. Table III provides the two-way Wilkoxon ranksum statistic test for several comparisons of the three main metrics.

**TABLE II.**
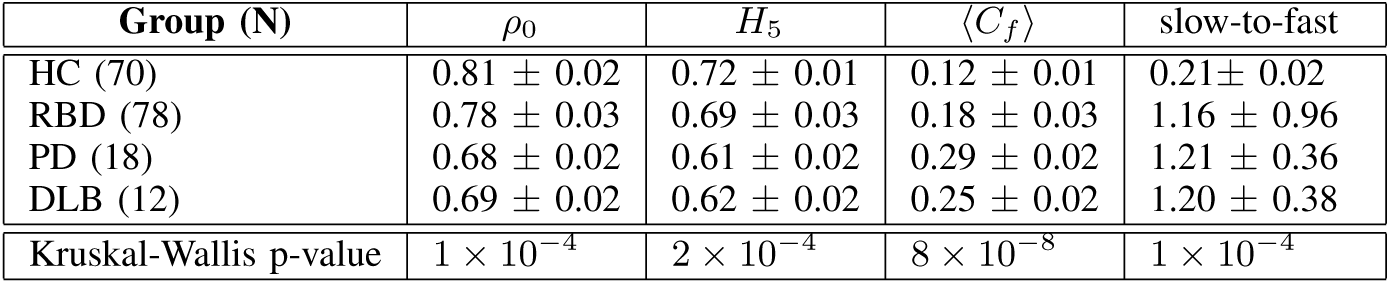
MEAN COMPLEXITY (*ρ*0), ENTROPY RATE (*H*5), FREQUENCY MUTUAL ALGORITHMIC INFORMATION MEAN CLUSTERING COEFFICIENT ⟨ *Cf* ⟩ (FOR A THRESHOLD OF 0.0425) AND SLOW-TO-FAST METRICS FOR EACH GROUP, WITH STANDARD ERROR OF THE MEAN (SEM). IN THE LAST ROW, THE FOUR-GROUP KRUSKAL-WALLIS P-VALUE FOR EACH METRIC IS PROVIDED.

**TABLE III.** STATISTICS FOR COMPARISONS BETWEEN GROUPS FOR SELECTED METRICS: WILKOXON RANK-SUM STATISTIC SIGNIFICANCE TWO SIDED P-VALUE AND THE AREA UNDER THE CURVE (AUC, IN PARENTHESIS).

| Comparison | $\rho_0$ | $\langle C_f \rangle$ | slow-to-fast |
| --- | --- | --- | --- |
| HC vs. RBD | $1 \times 10^{-1}$ (57%) | $3 \times 10^{-3}$ (63%) | $5 \times 10^{-3}$ (63%) |
| HC vs. PD | $1 \times 10^{-4}$ (**80%**) | $7 \times 10^{-7}$ (**88%**) | $2 \times 10^{-2}$ (69%) |
| HC vs. DLB | $3 \times 10^{-3}$ (**77%**) | $7 \times 10^{-4}$ (**82%**) | $9 \times 10^{-5}$ (**85%**) |
| RBD vs. PD | $2 \times 10^{-2}$ (73%) | $1 \times 10^{-4}$ (**80%**) | $5 \times 10^{-1}$ (55%) |
| RBD vs. DLB | $3 \times 10^{-2}$ (69%) | $4 \times 10^{-2}$ (70%) | $8 \times 10^{-3}$ (74%) |
| PD vs. DLB | $6 \times 10^{-1}$ (56%) | $3 \times 10^{-1}$ (62%) | $4 \times 10^{-1}$ (59%) |
| HC vs. PD+DLB | $7 \times 10^{-6}$ (**78%**) | $2 \times 10^{-8}$ (**86%**) | $6 \times 10^{-5}$ (**75%**) |
| HC vs. rest | $3 \times 10^{-3}$ (63%) | $6 \times 10^{-6}$ (69%) | $2 \times 10^{-4}$ (67%) |
| HC+RBD vs. PD+DLB | $2 \times 10^{-5}$ (**75%**) | $2 \times 10^{-7}$ (**80%**) | $1 \times 10^{-3}$ (69%) |
| RBD vs. PD+DLB | $5 \times 10^{-4}$ (**72%**) | $4 \times 10^{-5}$ (**76%**) | $5 \times 10^{-2}$ (62%) |

**Fig. 3.**
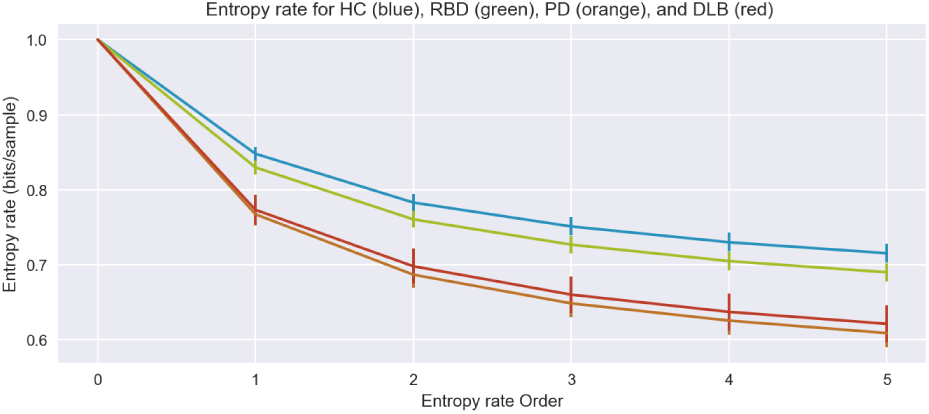
Entropy rate *H*_*n*_ as a function of order. Entropy rate plot with error bar (standard error of the mean) as a function of order.

**Fig. 4.**
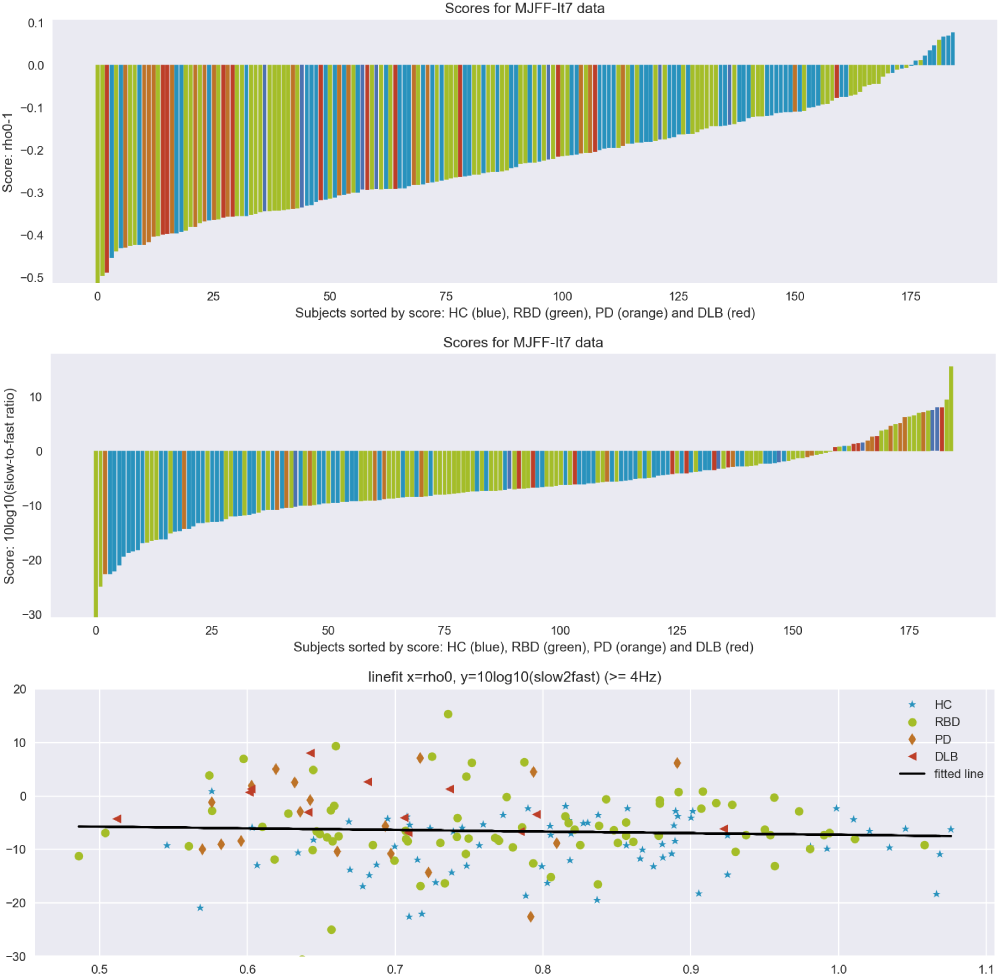
Complexity and (log) slow-to-fast scores per subject using full-band spectrograms. Complexity metric (*ρ*_0_ *-* 1 is shown for convenience) per subject for each class (HCs in blue, RBDs in green, PDs in orange and DLBs in red). The second one displays the log of slow-to-fast metric and the last displays *ρ*_0_ vs log of the slow-to-fast (line fit: slope: slope:-3.03 *±* 0.06, r: −0.07, p-value: *p <* 0.5).

**Fig. 5.**
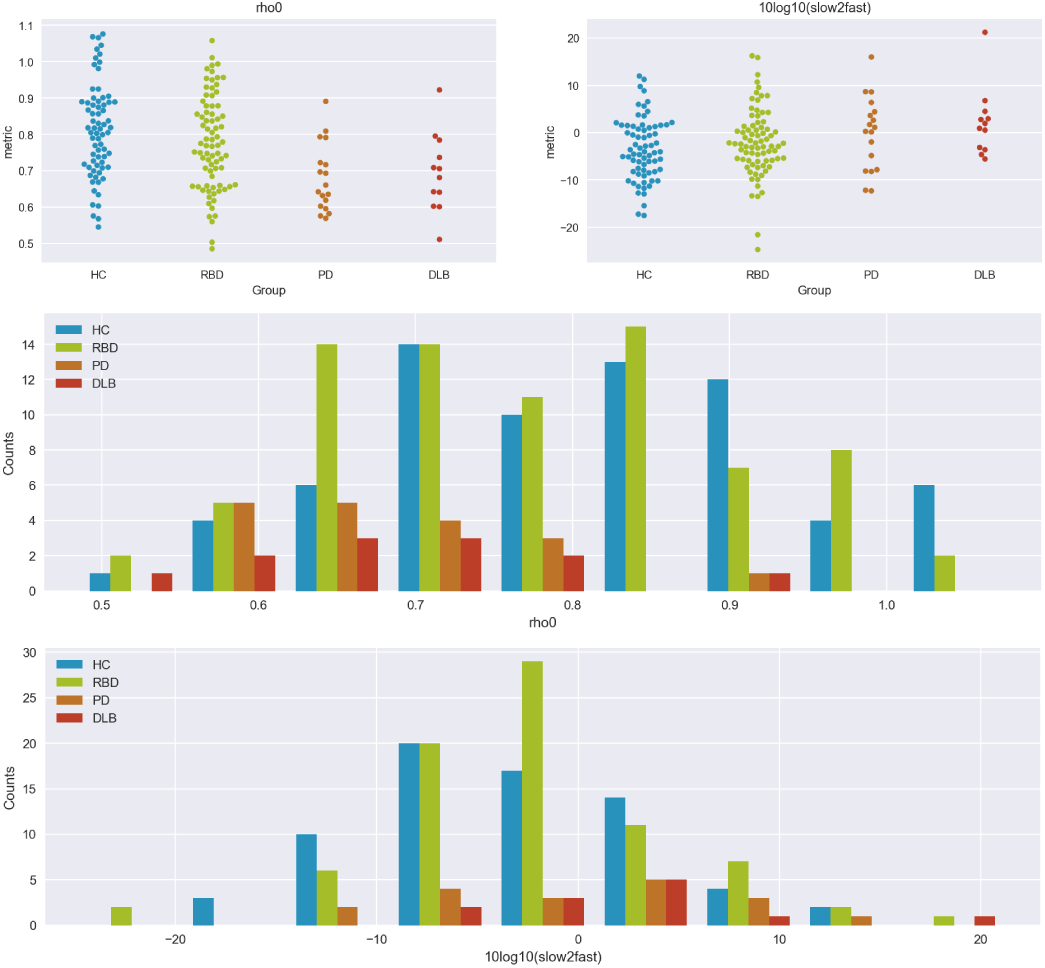
Histograms and scatter plots for *ρ*_0_ complexity and slow-to-fast metrics from full-band spectrograms. Scatter plots (top), and histograms for *ρ*_0_ and slow-to-fast with each class (HCs in blue, RBDs in green, PDs in orange and DLBs in red).

For completeness, we explored complexity applied on an epoch by epoch basis, which allowed us to use all the data from every subject (string length prior compression not being dependent on number of epochs). Complexity per epoch was averaged, leading to an overall increase in complexity but slightly improved statistical performance (*p <* 3 ×10^*-*5^ four group Kruskal-Wallis).

For comparison, we have also computed the *slow-to-fast* frequency power ratio, normally defined as [(*δ* + *θ*)*/*(*α* + *β*)], with *δ* [0.5,4 Hz), *θ* [4,8 Hz), *α* [8,13Hz), and *β* [13,32Hz) [7]. Here we computed it as the ratio of power in the interval [4, 8) Hz (*θ*) frequencies versus [8, 32) Hz (*α* +*β*), which we saw gave much better performance than the canonical one (which produced barely significant four-group p*<*0.05 Kruskal-Wallis statistical test results).

Finally, we studied the impact of using a few channels instead of the full set of 14. Complexity metrics remained robust, indicating that relevant information for statistical discrimination is available in each channel. For example, using *ρ*_0_ with stacks generated from channels P4 and P7 led to a statistically significant results in the four-group Kruskal-Wallis analysis (*p <* 2 × 10^−4^). The same analysis using the slow-to-fast ratio metric was not significant.

### A. The role of frequency in complexity and compressibility

To better understand changes in compressibility in PD and DLB converter subject group data, we investigated the role of high versus low frequency bands in complexity. Shortly, if only low frequency data (*f <* 14 Hz) is used to produce binarized strings, complexity metrics fail to discriminate the subject classes well. Similarly, restricting binarization and analysis to either the *β* or *γ* band alone eliminated statistical differences in *ρ*_0_ across the groups. If only high frequency data is retained (*f >* 12 Hz), statistical performance (separability) was partially maintained (*p <* 2 × 10^−2^ four-group Kruskal-Wallis).

Next, using the concept of mutual algorithmic information described above, we studied differences across groups from graphs constructed using MAI across frequency bands (*μ*_0_) using complex network theory [29], reasoning that there may be an increase in mutual information *across* bands that partly explains changes in compressibility of spectrograms (as discussed further below). Table II provides the main statistics for the mean frequency clustering coefficient—the clustering coefficient ⟨*C*_*f*_⟩ derived from the mutual algorithmic information—, and Figure 6 provides the degree and clustering coefficient as a function of threshold and the associated graph for a representative threshold.

**Fig. 6.**
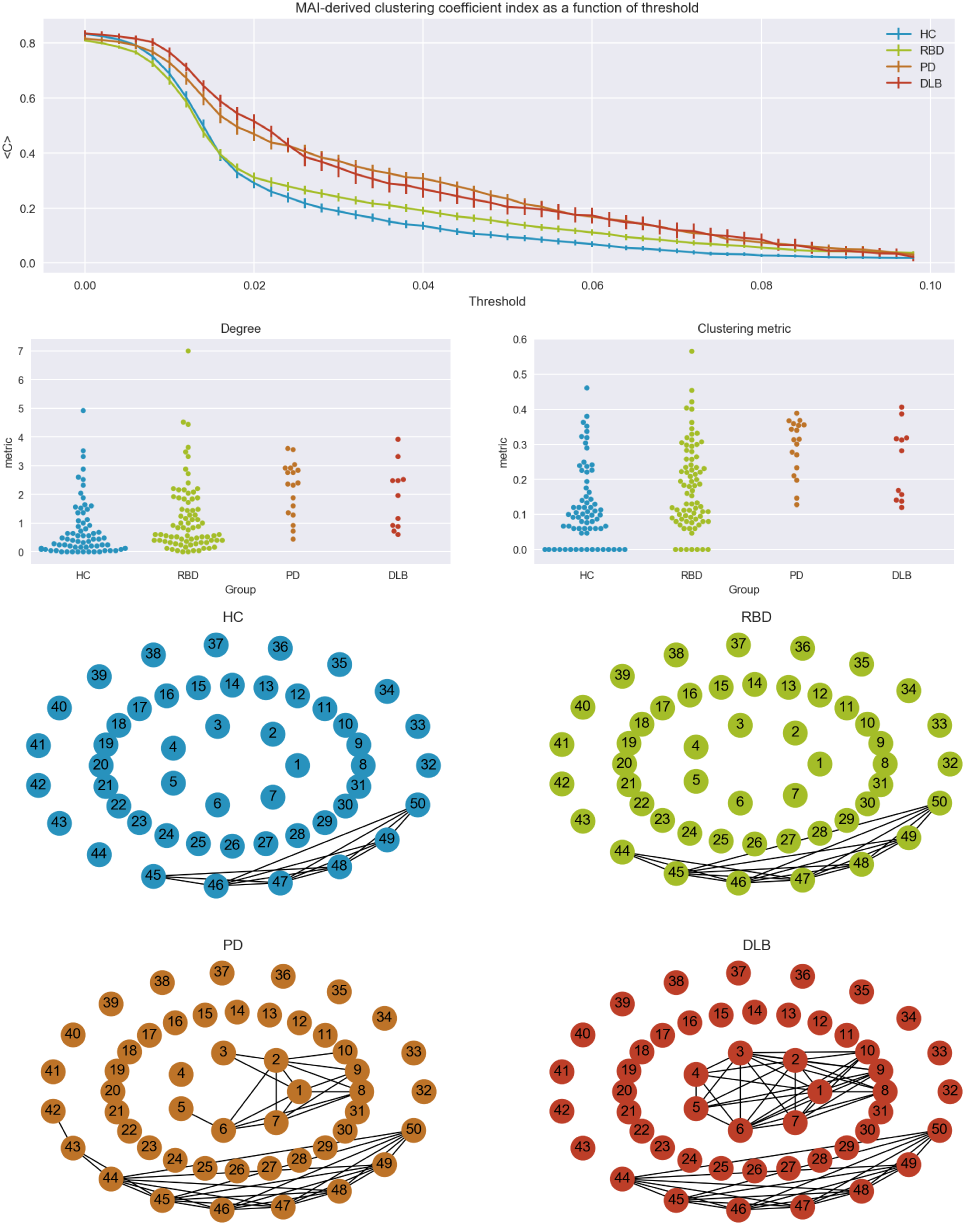
Graph analysis using connectivity from algorithmic mutual information (MAI, *μ*_0_) across frequency bands. Top: clustering coefficient as a function of threshold for the different groups. Middle: corresponding scatter plots. Bottom. Average graphs (threshold = 0.0425), with each node representing a frequency bin (label in Hz). Note the increasing MAI within low and within high bands in the second supergroup (PD+DLB).

## IV. DISCUSSION

Using a few minutes of eyes-closed resting EEG data to generate spectrogram stacks, we have seen here that RBD patients who later developed PD or DLB display diminished complexity compared to HCs or to RBD patients who remained disease-free. The results of our statistical analysis are compelling, displaying significant differences between two main super-groups: HCs and RBDs vs. PDs and DLBs. Data from the latter group display lower complexity, entropy rates and increased cross-frequency amplitude coupling. We have also seen that as expected, LZW and entropy rate provide very similar metrics (although LZW is much faster to compute).

These differences are remarkable as they reflect brain state at the moment of idiopathic RBD diagnosis, years before conversion to PD or DLB. The performance of these metrics is slightly superior and complementary to those from power analysis [6], [7], with which they are well aligned. Unlike in previous studies in neurodegenerative diseases [19], [21], [31], power ratio and complexity metrics were not correlated here (we note that correlation of power metrics such as slow-to-fast ratio and LZW or entropic complexity would not be entirely surprising, since complexity analysis is based on compression of spectrograms and EEG power imbalances that can lead to more compressible spectrograms after binarization). In addition to improving statistical separability, we note that the global complexity metric provides a rather general detection mechanism of regularities in the data. For example, imbalances in power ratio across arbitrary different bands can lead to lower complexity after data binarization. Complexity metrics can detect such regularities in the data without the need to define a priori which specific aspects to use (e.g., which bands to use for a ratio computation). In this sense, they provide a rather assumption-free analysis of the data which can then be further dissected. Moreover, they remained robust even if applied (and then averaged) epoch by epoch or using a few channels, which could be useful for screening in clinical practice.

### A. Analysis of sources of compressibility

As suggested above, even if complexity metrics detect differences across groups, it is interesting to investigate where specifically the relevant regularities lie in the data. They may lie in the temporal series within bands and across epochs, similarities across bands or across channels—that is, in frequency, time and space. Where is the loss of complexity taking place in patients with poor prognosis?

To find out in detail, we carried out tests reordering and randomizing the data in different ways to detect which lead to a loss of discrimination between the groups using complexity metrics (see Figure 8 for a visual representation of the different data re-shuffling methods). In general, all such tests led to an overall increase in complexity, highlighting the fact that compression exploits regularities across channels, across time (within and across frames), across frequency and across epochs, and that by randomizing the data it becomes harder to compress (more complex). However, interestingly, not all the above reshuffling methods led to a loss of discrimination across groups.

**Fig. 8.**
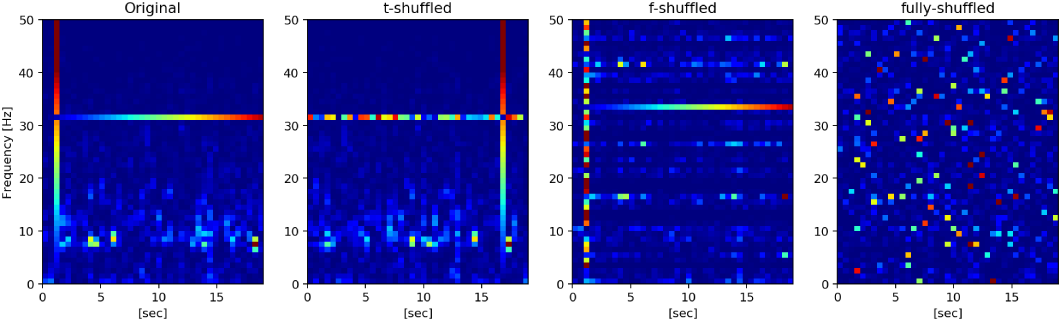
Frame (intra-epoch) spectrogram shuffling. From left to right: original, temporally shuffled, frequency and fully shuffled spectrograms. A horizontal and a vertical line has been added for reference.

We first randomized the frequency index at each time point within each frame independently across frames. As expected, this shuffling destroyed the discrimination performance of both complexity and power ratio metrics. However, if the frequency randomization was held constant within each spectrogram frame (but randomized across frames), only the performance of the power ratio metric (slow-to-fast) was severely affected. Similarly, applying the same randomization procedure across the temporal dimension (reshuffling time within each frame, which leaves power ratio metrics unaffected, as they rely on unordered time averages) increased overall complexity but did not affect the discriminability of the complexity metric. Thus, complexity metrics appear to rely on within-frame temporal regularities across some frequencies (similar patterns appearing in several frequencies or at several time points in a frame), with highest complexity for HC and RBD, then PD and finally DLB—irrespective of the actual shape of the spectrum. Poor prognosis in RBD appears to be associated with a decrease in cross-frequency, within-frame temporal regularities in spectrograms. A partial explanation of the above rests on the observation that binarization by the median of the global stack can lead to simple sequences in some frequencies bands which naturally tend to be far away from the global median, i.e., low and high frequencies, which have power typically either above or below the median. Frequency bands whose values lie far from the median will tend to look simpler (producing mostly chains of 0’s or 1’s). Thus, our complexity metric applied to global stacks in which either frequencies or channels lie far from the median produces lower complexity estimates. The complexity metric is insensitive as to where these regularities are (in which frequency or channel)—it detects events of similar symbol aggregation—chains of 0’s or of 1’s.

A further test we carried out was to binarize each band independently, which in essence removes differences in power scale across bands since binarization then uses each band’s median. This resulted in a loss of discriminability using *ρ*_0_, indicating that imbalances in power across bands are an important element in *ρ*_0_ complexity. However, as discussed above, we observed that cross-frequeny MAI differences across groups remained significant and discriminative despite this manipulation (see, as an example, Figure 7). This implies that there exist scale-independent cross-frequency (amplitude) temporal regularities in the data (typically an increase in regularity with disease progression).

**Fig. 7.**
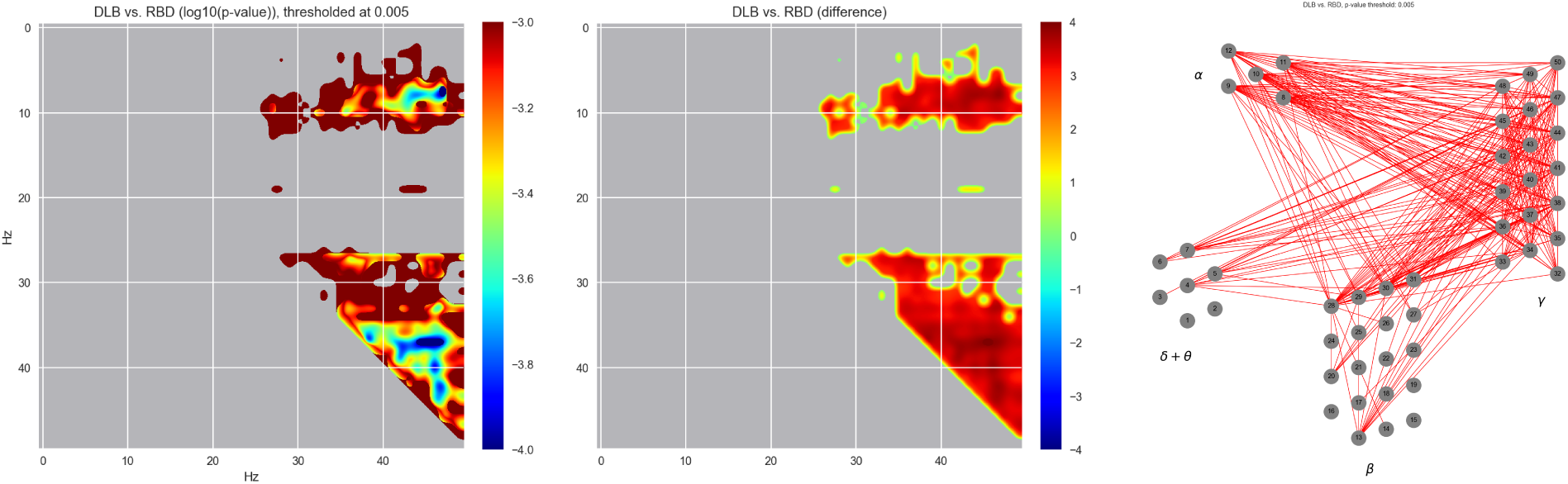
Statistics of MAI graphs (*μ*_0_) across frequency bands. Left: statistical differences between two groups at the edge level using per band binarization to remove amplitude scale differences. Right. Associated graph displaying links with increased (red) or decreased (blue) scale-independent MAI connectivity in the first group with respect to the second. In this particular case, we observe an increase in connectivity, especially in alpha+theta to gamma bands.

Our results with regard to the role of high frequencies in changes in complexity can be related to those in [31], where multiscale entropy was employed as complexity metric to study the differences in brain signal variability in PD patients who developed dementia at follow-up with respect who did not. The authors found significant changes, with lower signal variability at timescales sensitive to higher frequencies (i.e., more compressibility at higher frequencies) in patients that developed dementia than those who did not or in controls. We do note, however, that multiscale entropy and LZW (or entropy rate) estimate complexity in different ways.

Finally, one can ask how much does stack compression benefit from having access to all frames from a subject as opposed to compressing each frame independently. The answer is that PD and DLB data benefit the most (with a ratio of per frame vs. per stack *ρ*_0_ of 1.26). That is, using the full stack data improves compression by 26% in these groups, compared to 22 and 23% in HC and RBDs (*±*1%, p *<*6 × 10^−5^ four group Kruskal-Wallis). There are longer term persistent regularities in the PD and DLB groups than in HC or RBD.

## V. CONCLUSION

In [11] it is hypothesized that while a system capable of universal computation may produce many different types of patterns, both simple (e.g., constant or repetitive) and complex, a healthy brain engaging in modeling and prediction of complex input-output strings will produce complex-looking, highly entropic data. Such apparent complexity is what we actually measure when using entropy rate and LZW compression of electrophysiological or metabolic brain data [13]–[17]. First order entropy, entropy rate or LZW provide (poor) *upper bounds* on algorithmic complexity.

From our results with this dataset we conclude that LZW and, equivalently, entropy rate highlight losses of estimated algorithmic complexity in spectrograms of RBD patients likely to evolve into PD or DLB compared to those that remained in RBD or to healthy controls. Our full band complexity scores are generally more discriminative than spectral ratios, do not require a pre-selection of the spectral bands to study, and remain robust even when using a few channels. The loss of complexity takes place both in low and high frequencies, and is partly due to increased mutual information within and across bands. These results indicate that information *differentiation* (global complexity) is a potentially relevant metric for the prognosis of RBD, and are in line with current views of the brain stemming from information theory connecting theories of cognition and consciousness with the phenomenology of brain health, and in particular, neurodegeneration [11], [32].

## AUTHOR CONTRIBUTIONS

DI and MC pre-processed EEG data to produce artifact free spectrograms. EK and ASF contributed to code development and revised the manuscript. JFG, RP and JM collected the EEG data and revised the manuscript. GR provided the complexity analysis code, carried out the complexity and statistical analysis and wrote the manuscript.

## ACKNOWLEDGMENTS

This work partly supported by the Michael J. Fox Foundation project “Discovery of EEG biomarkers for Parkinson Disease and Lewy Body Dementia using advanced Machine Learning techniques” (Rapid Response Innovation Awards 2013), the FET Open Luminous project (H2020-FETOPEN-2014-2015-RIA under agreement No. 686764) as part of the European Union’s Horizon 2020 research and training programme 2014-2018, the Canadian Institutes of Health Research (CIHR) and the W. Garfield Weston Foundation.

**Giulio Ruffini** Giulio graduated (math/physics) from UC Berkeley and obtained a PhD in physics from UC Davis (1995). In 2000, he co-founded Starlab to transform research into technologies with positive impact. During the FET HIVE project, his team developed multielectrode hybrid EEG-transcranial stimulation devices, and he recently led the first demonstration of noninvasive brain-to-brain communication. In 2011, he co-founded Neuroelectrics to deliver clinical EEG-tCS systems. He collaborates with teams worldwide developing applications of EEG and noninvasive stimulation from the integration of biophysics, computational models and AI, as in the FET Open Luminous project studying consciousness.

